# Indirect genetic effects across ontogeny in an avian cooperative breeder

**DOI:** 10.64898/2026.05.16.725675

**Authors:** Gladiana Spitz, David Tian, Elissa Cosgrove, Tori D. Bakley, Sahas Barve, Reed Bowman, John W. Fitzpatrick, Nancy Chen

## Abstract

Social interactions are ubiquitous in nature and have the potential to affect trait evolution, particularly in group-living animals such as cooperative breeders. Interactions among conspecific individuals can affect the amount of additive genetic variation for a trait when the phenotype of an individual is also affected by the genotype of its social partner(s) via indirect genetic effects. Thus, quantifying both direct and indirect genetic effects of social partners is critical for understanding and predicting evolutionary trajectories. While much is known about maternal indirect genetic effects, empirical estimates of indirect genetic effects from other social partners remain limited, particularly in wild populations. Here, we use animal models to assess the contribution of indirect genetic effects from all social partners in a family group (mothers, fathers, and helpers) on juvenile morphometric traits across ontogeny in the cooperatively-breeding Florida scrub-jay (*Aphelocoma coerulescens*). We found indirect genetic effects of helpers and fathers on nestling weight, but no indirect genetic effect of mothers. Across ontogeny, we found increasing additive genetic variation in both weight and tarsus length. Our study provides a comprehensive assessment of within-group indirect genetic effects in a cooperative breeder and highlights the importance of considering indirect genetic effects beyond maternal effects.

## Introduction

Social interactions occur between conspecific individuals of all taxa and can play an important role in phenotypic evolution in nature (Cheverud and Moore 1994; Moore et al. 1997; Wolf et al. 1998). An individual’s phenotype may be influenced by both the environment experienced by a social partner (indirect environmental effects) and the genotypes of a social partner (indirect genetic effects) (Moore et al. 1997; Wolf et al. 1998). A recent meta-analysis found that on average, accounting for indirect genetic effects leads to a 66% increase in the total amount of genetic variation available to selection, relative to narrow-sense heritability (Santostefano et al. 2025). Indirect genetic effects have the potential to change the evolutionary trajectories of phenotypes by amplifying, dampening, or even reversing a phenotype’s response to selection (Wolf et al. 1998; Bijma and Wade 2008; McGlothlin and Brodie III 2009). Not accounting for indirect genetic effects can also inflate estimates of narrow-sense heritability (*h*^*2*^), leading to overestimated responses to selection (Kruuk 2004). Thus, quantifying both direct and indirect genetic effects of social partners is critical for understanding and predicting evolutionary trajectories.

While the theoretical frameworks for estimating indirect genetic effects are well developed (Moore et al. 1997; McGlothlin and Brodie III 2009; Bijma 2010), empirical studies of indirect genetic effects in wild populations remain limited, as few systems have sufficient relatedness and phenotypic data to robustly estimate indirect genetic effects (Baud et al. 2022). In addition, many studies have thus far focused solely on maternal effects (McAdam et al. 2002, 2014; Wilson and Réale 2006; Gauzere et al. 2020) or the effect of competitors (Bleakley and III Brodie 2009; Wilson et al. 2010; Fisher et al. 2019; Santostefano et al. 2021), and not yet considered the full spectrum of potential social interactions that an individual may experience.Paternal interactions are often considered to be purely environmental or mediated by maternal effects, but a growing body of literature suggests paternal effects as important in their own right (Crean and Bonduriansky 2014). For example, in dung beetles (*Onthophagus taurus*), males with larger horns increase brood mass by helping the females with offspring provisioning (Hunt and Simmons 2000). Indirect genetic effects may also go beyond social interactions between parents and offspring, particularly in cooperatively breeding systems where individuals live in groups composed of breeding parents and non-breeding helpers (Skutch 1935). Helpers are mature but non-breeding individuals who contribute to territory defense, offspring care, and resource acquisition (Skutch 1935). As a result, helpers may confer indirect genetic effects. Indirect genetic effect models that incorporate multiple individuals necessitate large sample sizes (Bijma 2014; Fisher et al. 2019), and thus far few empirical studies have estimated the indirect genetic effect of helpers (but see Adams et al. (2015) for helper effects on parents). By not considering all possible social interactions, we lack a complete understanding of trait evolution and evolutionary potential.

The presence and strength of indirect genetic effects may not only vary across individuals, but also across ontogeny. The strength of caregiver effects is expected to decrease as individual offspring age, as direct interactions with parents tend to decrease as offspring age (Wilson and Réale 2006; Pick et al. 2016; White and Wilson 2019; Gauzere et al. 2020). For example, maternal effects in domestic animals (*Bos* and *Ovis*) and red deer (*Cervus elaphus*) as well as paternal effects in burying beetle (*Nicrophorus vespilloides*) have been found to decrease as offspring age (Wilson and Réale 2006; Head et al. 2012; Gauzere et al. 2020). Whether helper effects on juvenile traits similarly decrease across ontogeny remains uncertain. Nevertheless, measuring how indirect genetic effects may vary as offspring develop is important, as it allows us to identify when social interactions most strongly influence total heritable variation and evolutionary trajectories.

Here, we leverage a long-term study of the Federally threatened Florida scrub-jay (*Aphelocoma coerulescens*) population at Archbold Biological Station (Woolfenden and Fitzpatrick 1984) to assess indirect genetic effects of different social partners on offspring morphology across ontogeny. Florida scrub-jays perform biparental care and are cooperative breeders; most individuals delay dispersal to assist their parents in rearing offspring on their natal territories (Woolfenden and Fitzpatrick 1984). During the nestling stage, the male breeder and any helpers present on the territory perform the majority of food provisioning of offspring while the female breeder incubates (Woolfenden and Fitzpatrick 1984). Once the offspring have fledged, both parents and helpers assist in food provisioning until the juveniles reach nutritional independence (Woolfenden and Fitzpatrick 1984). While helpers have a beneficial effect on nestling weight, they can become competitors for food once offspring reach nutritional independence (Woolfenden and Fitzpatrick 1984; Mumme et al. 2015).

We characterized maternal, paternal, and helper effects on offspring weight and size at two developmental life stages using animal models. Offspring weight is positively associated with survival (Mumme et al. 2015), and decomposing variation in offspring weight might provide insight into additional factors that could affect survival. To disentangle the effects of these different social partners within a family group, we fitted models that included single and multiple social effects. Based on differences in food provisioning behavior, we expected fathers and helpers to contribute towards nestling traits more than mothers. Additionally, we expected a shift toward stronger direct genetic effects and weaker indirect genetic effects as juveniles reach nutritional independence. This study investigates indirect genetic effects from an entire caregiving unit on offspring morphometric traits at different life stages, providing new insights into the drivers of phenotypic evolution.

## Materials and Methods Study Species and Population

The Florida scrub-jay is a cooperatively breeding and highly-territorial bird restricted to xeric oak scrub in central Florida (Woolfenden and Fitzpatrick 1984). Family groups consist of the monogamous breeding pair, their juvenile offspring, and up to seven helpers (Woolfenden and Fitzpatrick 1984). While Florida scrub-jays do form long-term pair bonds, individuals will pair with another jay when widowed or in cases of divorce, so many individuals end up with multiple partners over their lifetime (Woolfenden and Fitzpatrick 1984). Helpers are typically non-breeding adult offspring of the social breeders, though individuals sometimes disperse to join unrelated groups(Woolfenden 1975; Suh et al. 2022). Florida scrub-jay nestlings fledge or leave the nest around 15-20 days after hatching, and fledglings attain nutritional independence around 60-90 days after hatching (Woolfenden and Fitzpatrick 1984; Mumme et al. 2015).

We used data from an intensive, long-term study of a population of Florida scrub-jays at Archbold Biological Station in Highlands County, Florida (Woolfenden and Fitzpatrick 1984). All individuals are uniquely color-banded, and the entire population is censused once a month. The territory of each family group is mapped each year. Nearly all nests are located during building, laying, or incubation and then closely monitored to collect accurate data on clutch sizes and hatching dates. Nestlings are banded, sampled for blood, and measured 11 days after hatching and again 50-100 days after hatching. Here, we only considered the years for which we have blood samples and molecular sexing data for >70% of the birth cohort: 1989-1991, 1995, and 1999-2021. We excluded the small number of nests with known extra-pair paternity. Our final dataset included 3,313 individuals from 185 territories. All work was approved by the Cornell University Institutional Animal Care and Use Committee (IACUC 2010-0015) and authorized by permits from the US Fish and Wildlife Service (TE824723-8), the US Geological Survey (banding permit 07732), and the Florida Fish and Wildlife Conservation Commission (LSSC-10-00205).

## Relatedness Information

Extra-pair paternity in the Florida scrub-jay is rare (Quinn et al. 1999; Townsend et al.2011), thus we can construct accurate pedigrees from field observations alone. We verified relationships in the pedigree using genomic data collected for 3,583 individuals in 1989–1991, 1995, and 1999–2013 (Chen et al. 2016). We pruned the pedigree to only include ancestors of individuals in our dataset using the prunePed function from the R package MCMCglmm (Hadfield 2010). Our pruned pedigree had 4,574 records, with 4,070 parent-offspring trios, a mean pedigree depth of 5.7 generations, and a maximum depth of 14 generations.

### Quantitative Traits

We analyzed individual weight and tarsus length (as a proxy for size) across two different developmental stages to explore the magnitude of indirect genetic effects over time. We measured nestlings on day 11 after hatching and juveniles between 60-90 days after hatching (Table 1). Both nestling and juvenile weight are significant predictors of postfledging survival (Mumme et al. 2015).

**Table 1.** Summary statistics for each trait included in this study. *n* is the sample size, mean is the mean trait value, and CV is the coefficient of variation, calculated as standard deviation/mean.

| Trait | $n$ | Mean | CV |
| --- | --- | --- | --- |
| Nestling Weight (g) | 3198 | 43.77 | 0.16 |
| Juvenile Weight (g) | 1353 | 69.86 | 0.065 |
| Nesting Tarsus (mm) | 2039 | 29.37 | 0.087 |
| Juvenile Tarsus (mm) | 1341 | 36.32 | 0.032 |

### Variance partitioning model

To partition the total phenotypic variance of each trait into direct and indirect genetic and environmental components, we fitted univariate linear mixed-effect models known as animal models with a relatedness matrix calculated from the pedigree (Falconer and MacKay 1996; Kruuk 2004; Wilson et al. 2010). We used a similar model as Fisher et al. (2019) and Santostefano et al. (2021):

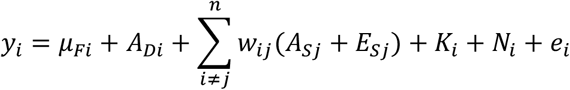

where *y*_*i*_ is the phenotype of individual *i*, μ_*Fi*_ is the population mean adjusted for fixed effects, *A*_*Di*_ is the direct additive genetic effect, *W*_*ij*_ is a parameter specifying how indirect effects scale with group size, *A*_*sj*_ is the indirect additive genetic effect of social partner *j, E*_*sj*_ is the indirect environmental effect of social partner *j, K*_*i*_ is the natal year of individual *i, N*_*i*_ is the natal nest of individual *i*, and *e*_*i*_ is the residual.

To account for known sources of variation in offspring weight and tarsus length, we included the following fixed effects in all our models: sex, hatching date (day of year), brood size, and individual inbreeding coefficient. Hatching date, brood size, and inbreeding coefficient are negatively correlated with offspring weight in our study population (Mumme et al. 2015; Chen et al. 2016). We calculated individual inbreeding coefficients from the pedigree with the calcInbreeding function in the pedigree package (*v*. 1.4.2) (Coster 2022). As the number of helpers in a group is known to affect nestling and juvenile weight (Mumme et al. 2015), we included the number of helpers as a fixed effect when estimating maternal and paternal effects. To account for variation in age for juvenile measurements, we added age measured in days as an additional covariate to our juvenile models. Finally, as natal territory size has been shown to affect nestling and juvenile weight (Mumme et al. 2015), we included the size of the territory in hectares. We mean-centered all numeric fixed effects.

All models included random effects of individual ID to estimate additive genetic effects as well as natal year and identity of the natal nest to account for common environmental effects. To estimate maternal or paternal effects, we included the identity of the mother or father of the focal individual as a random effect. To estimate helper effects, we included the identities of each helper present at the natal territory of the focal individual as separate random effects. We assumed these random effects came from the same distribution with a mean of zero, allowing us to estimate a single variance term using a single design matrix (as in Fisher et al. (2019) and Santostefano et al. (2021)). As an individual’s indirect genetic effect may vary with group size, we weighted each helper random effect using the parameter *W*_*ij*_, calculated as 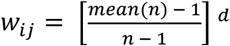, where *n* is the number of helpers + 3 and *d* is the dilution parameter (Bijma 2010). When *d* = 0, indirect genetic effects do not depend on group size, and when *d* = 1, indirect genetic effects are inversely proportional to group size (Bijma 2010). We determined the appropriate dilution parameter for each trait separately by fitting models using different values for *d* between 0 and 1, using intervals of 0.1. To test whether variation in relatedness between helpers and focal individuals affects helper contributions, we also fitted models with helper terms weighted by *r*_ij_, where *r*_ij_ is the pairwise relatedness between the focal offspring *i* and helper *j* calculated from the pedigree using the kinship function from the kinship2 package (*v* 1.9.6.1) (Sinnwell et al. 2024). We chose the value of *d* based on the model with the lowest Akaike Information Criterion (AIC; Supplemental Table 2).

### Model-fitting

For each trait, we fitted maternal, paternal, and helper effect models, as well as models with every combination of two or three types of social partners. We fitted all models using the restricted maximum likelihood approach implemented in ASReml-R 4.2 (Butler et al. 2023) in R 4.4.1 (R Core Team 2023). We tested the significance of each fixed effect using Wald F-statistic tests. To test the significance of random effects, we fitted a series of models, starting with a model with only fixed effects and residual variance (Model 1: V_R_), then sequentially adding random effects of natal year (Model 2: V_R_ + V_K_), natal nest (Model 3: V_R_ + V_K_ + V_N_), additive genetic effects (Model 4: V_R_ + V_K_ + V_N_ + V_DG_), social environment effects (Model 5: V_R_ + V_K_ + V_N_ + V_SE_), social genetic effects (Model 6: V_R_ + V_K_ + V_N_ + V_DG_ + V_SE_ + V_SG_), and the covariance between social genetic effects and additive genetic effects (Model 7: V_R_ + V_K_ + V_N_ + V_DG_ + V_SE_ + V_SG_ + cov(DG, SG)). We fitted models to estimate the effects of each type of social partner: mom (M), dad (D), or helper (H). As all the variance components were bound to be positive, we assumed an equal mixture of 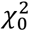and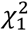when testing single variance components (Self and Liang 1987). As all three types of social partners interact within a single family group, we also fitted models with two social partners (M + D, M + H, D + H) or all three social partners (M + D + H). We compared models using AIC.

### Variance component calculations

We estimated the direct narrow-sense heritability (*h*^*2*^ = V_AD_/V_P_), indirect genetic effects 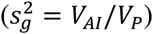,and indirect environmental effects 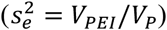by dividing the relevant variance component by the total phenotypic variance (V_P_). Total indirect effects were estimated by summing the indirect genetic and environmental effects: *s*^2^ = (*V*_*AI*_ + *V*_*pEI*_)/*V*_*p*_. We estimated the total heritable variation (V_H_) following Bijma (2010):

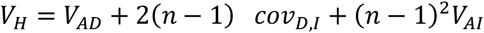

where cov_D,I_ is the covariance between direct and indirect genetic effects and *n* is group size. For models with a covariance structure that failed to converge, we set cov_D,I_ to 0. In the maternal and paternal effect models we set *n* as 2, while in the helper effect models we set *n* as the average number of helpers + 1. We estimated the relative total heritable variance *τ*^2^ = *V*_*H*_ /*V*_*p*_ as the total heritable variation divided by the phenotypic variance (Bijma 2010). Finally, we compared *τ*^2^ and h^2^ to assess the relative contribution of indirect genetic effects to total heritable variance (Bijma 2010). Differences between *τ*^2^ and *h*^2^ were assumed to be significant if the 95% confidence intervals did not overlap. We assumed estimates that were very small (<0.0001) were essentially zero.

## Results

### Additive Genetic Effects

We found significant additive genetic variance for all traits (comparison of Model 3 to 4; Table 2). The proportion of phenotypic variance explained by additive genetic effects (heritability, *h*^*2*^) varied among models for nestling weight, ranging from approximately zero for the model with all three social partners (Model M + D + H) to *h*^*2*^ = 0.14 for the model with direct genetic effects only (DGE-only; Model 4; Figure 1, Supp. Figure 1). Heritabilities for nestling tarsus length ranged from 0.046 to 0.083. In contrast, at the juvenile stage, heritabilities were similar across all models for both weight (*h*^*2*^ range = 0.43 – 0.47) and tarsus length (*h*^*2*^ range = 0.43 – 0.46) (Figure 2, Supp. Table 1). Across all models and traits, heritability was much greater at the juvenile stage than the nestling stage (Figure 1, Figure 2).

**Table 2:**
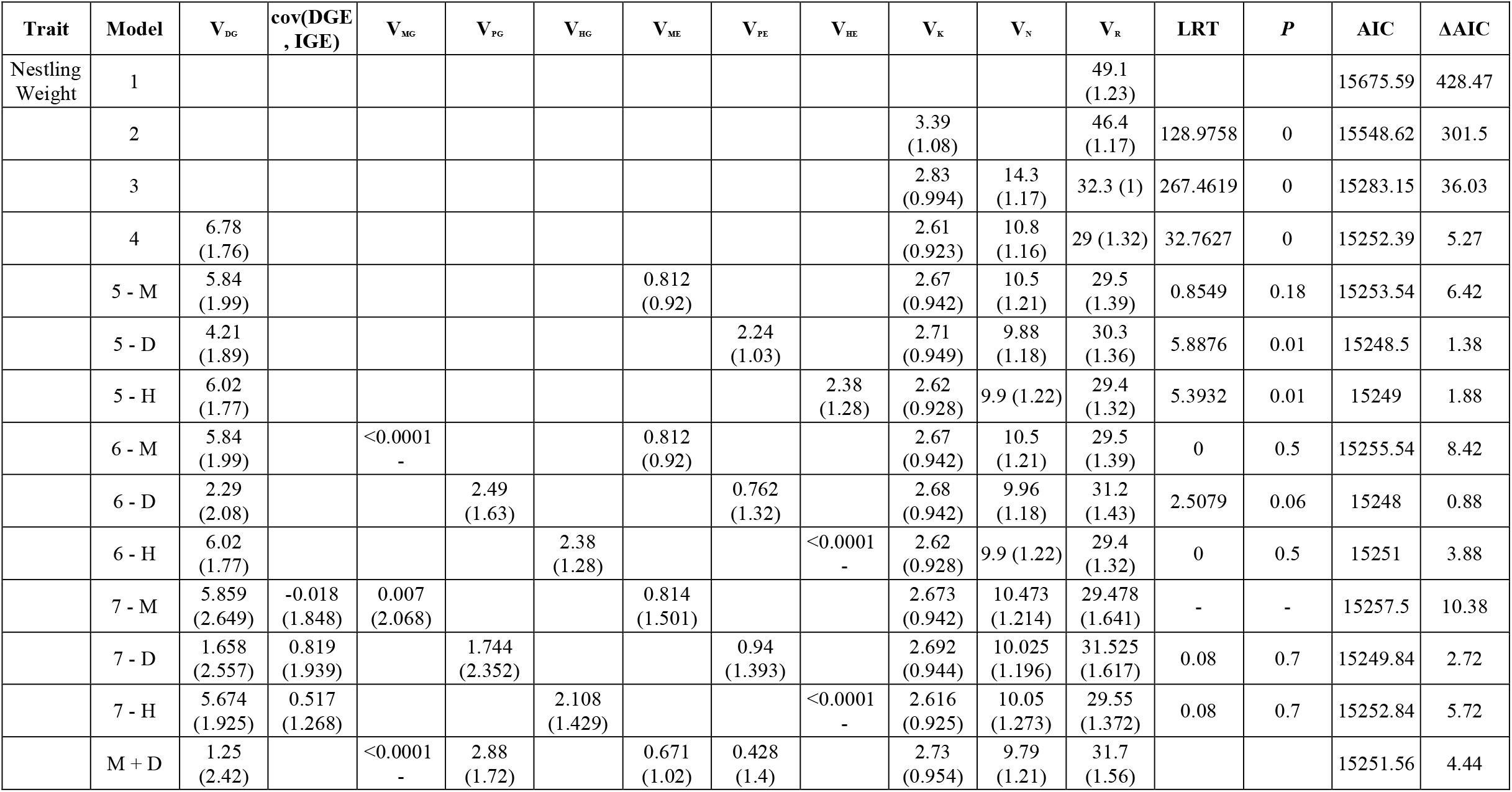
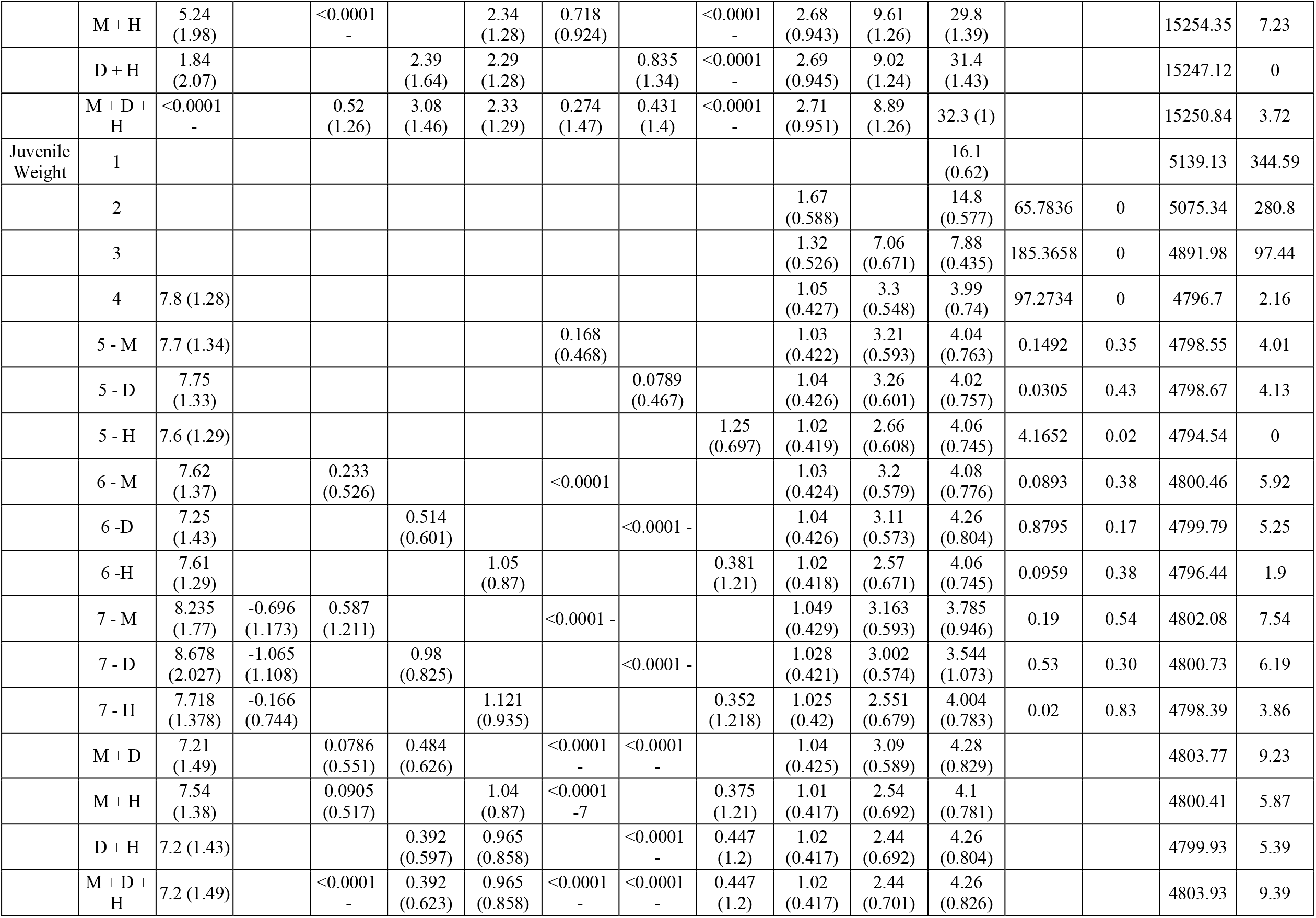
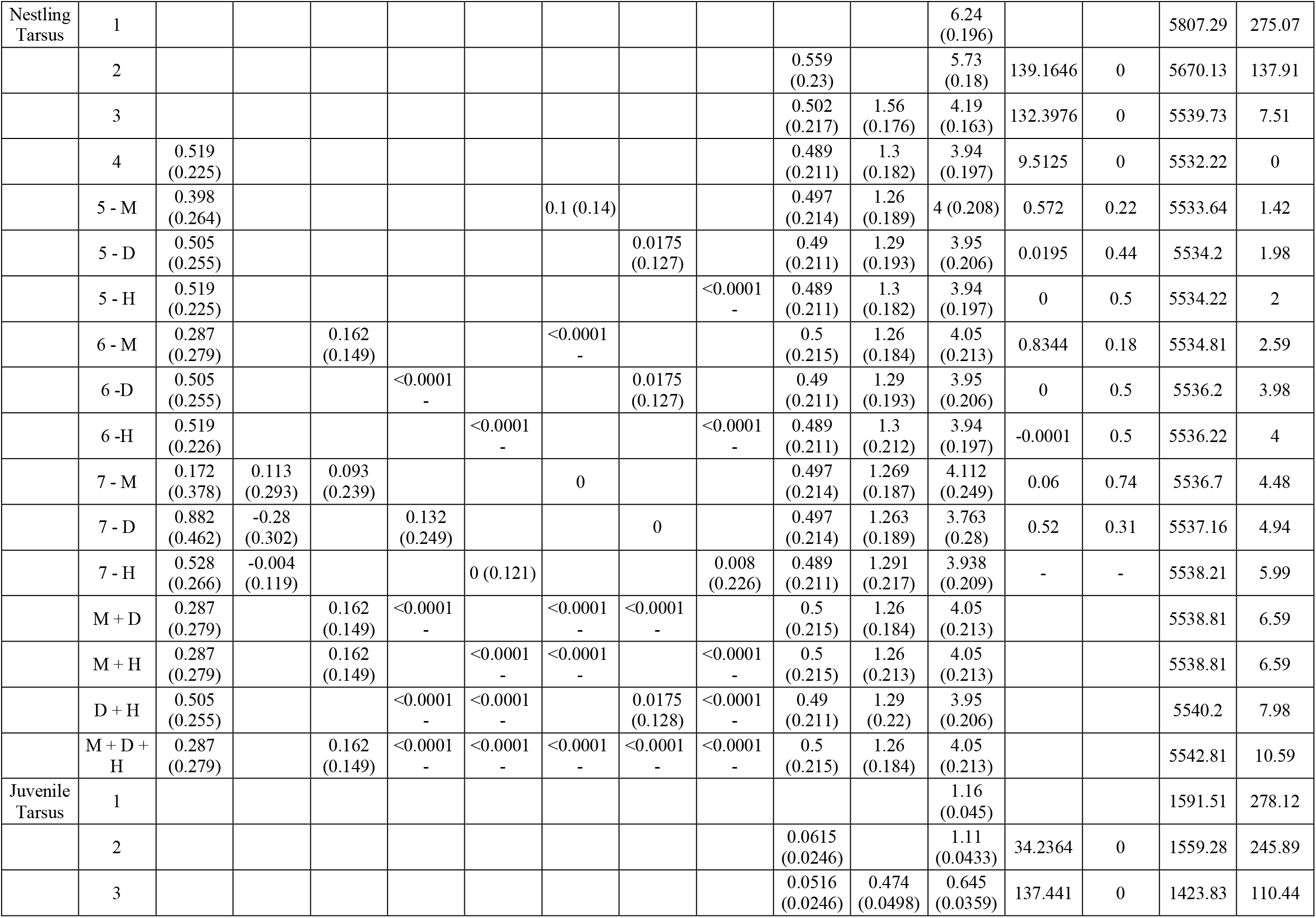
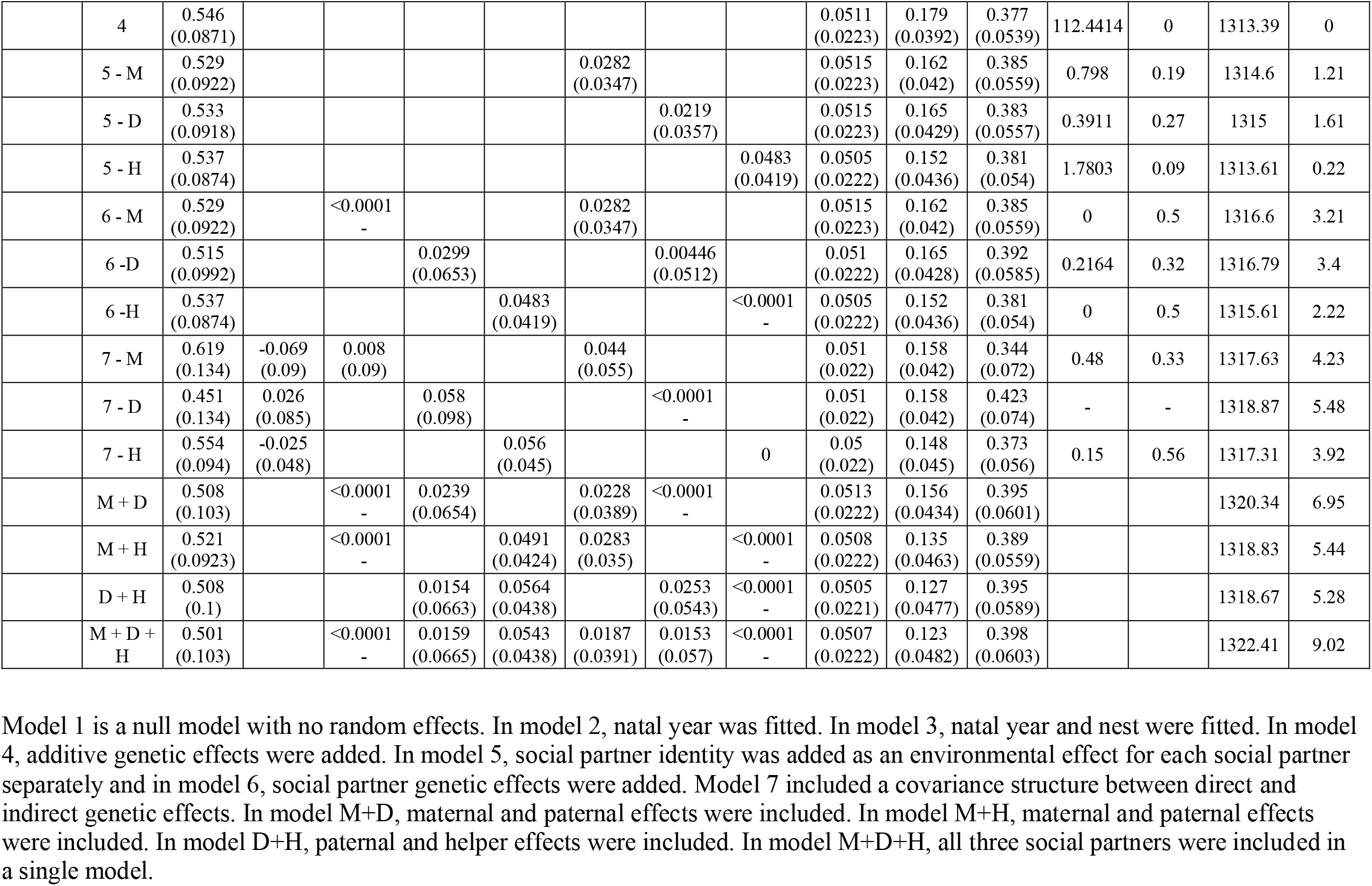
Variance components for all traits in each IGE model. Standard errors are in parentheses when available. V_DG_ is the direct genetic effect, V_MG_ is the maternal genetic effect, V_ME_ is the maternal environmental effect, V_PG_ is the paternal genetic effect, V_PE_ is the paternal environmental effect, V_HG_ is the helper genetic effect, V_HE_ is the helper environmental effect, V_N_ is the natal nest effect, V_K_ is the natal year effect, V_R_ is the residual, and cov(DGE,IGE) is covariance between direct genetic effects and indirect genetic effects. M = mom, D = dad, H = helper. We also show likelihood ratio test results between nested models and model AIC. ΔAIC is measured based on the best performing model. Three models that included a covariance structure between direct and indirect genetic effects did not converge, so we do not include the results of the LRTs for those models here.

**Figure 1:**
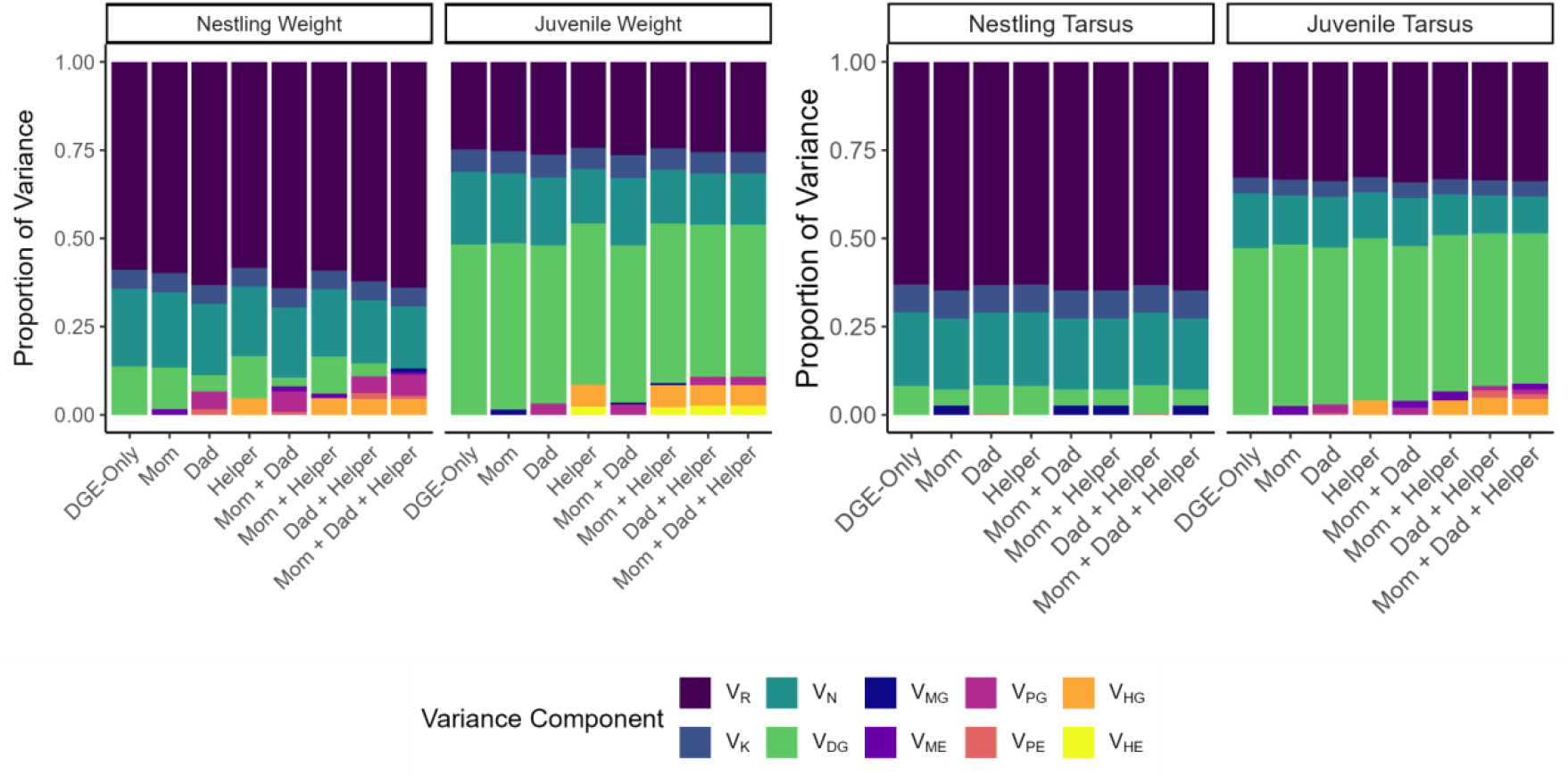
Estimated variance components as proportions of total phenotypic variance in offspring weight and tarsus length at the nestling and juvenile stages. For each trait, we ran DGE-only, maternal, paternal, and helper models, as well as models with multiple social partners to test the effect of different social partners. Colors represent different variance components. V_DG_ is the direct genetic effect, V_MG_ is the maternal genetic effect, V_ME_ is the maternal environmental effect, V_PG_ is the paternal genetic effect, V_PE_ is the paternal environmental effect, V_HG_ is the helper genetic effect, V_HE_ is the helper environmental effect, V_N_ is the natal nest effect, V_K_ is the natal year effect, and V_R_ is the residual. For full model results, see Table 2. Results including covariances between direct and indirect effects are shown in Supplementary Figure 1.

**Figure 2:**
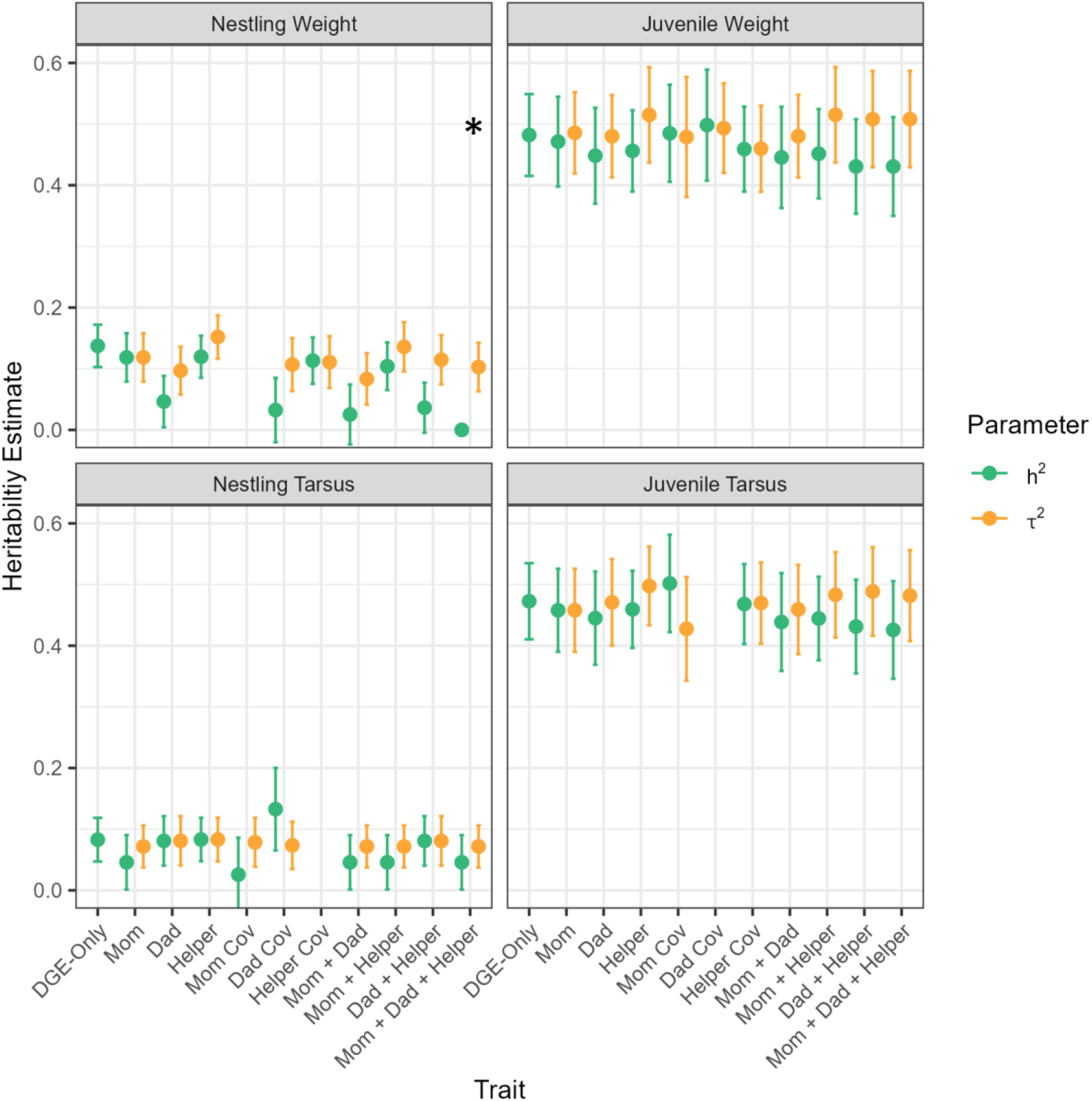
Narrow-sense heritability (*h*^*2*^) and relative total heritable variance (*τ*^*2*^) for nestling and juvenile weight and tarsus length for maternal, paternal, and helper models, as well as models with multiple social partners. Narrow-sense heritability and total heritable variance are similar across models for all traits except nestling weight. We do not show results for the three models that did not converge. Asterisks (*) denote a significant difference between *h*^*2*^ and *τ*^2^ based on 95% confidence intervals.

### Indirect Genetic and Indirect Environmental Effects

Dilution parameter testing found that weighting individual contributions of helpers by relatedness did not improve model fit, and the best dilution parameter was 1 for all traits except nestling tarsus, which did not depend on group size (Supp. Table 2, Supp. Table 3). A combination of paternal and helper genetic and environmental effects (Model D + H) was the best fit for nestling weight, though models with just paternal genetic effects (Model 6 - D), paternal environment effects (Model 5 - D), or helper environment effects (Model 5 - H) had equivalent support (∆AIC < 2; Table 2). In sum, indirect genetic effects from dads and helpers explained a significant amount of variation in nestling weight (*s*_*g*_^*2*^ = 0.093 ± 0.04) (Supp. Table 1). The best supported models for juvenile weight were models with helper environmental effects or helper genetic effects (Table 2). For tarsus length, the model with direct genetic effects (Model 4) had the highest support for both nestlings and juveniles, though including social environmental effects for any of the three social partners had equivalent support for juvenile tarsus length (Table 2). However, we did not detect significant indirect genetic or environmental effects for these three other traits (Supp. Table 1). Heritability and relative total heritable variance were not significantly different from each other, except for the model including all three social partners (Model M + D + H) for nestling weight (Fig. 2).

### Other Sources of Variance

Natal year and nest explained significant amounts of variance for both weight and tarsus length across all models (from comparisons of Model 1 and 2 as well as Model 2 and 3, respectively; Table 2). Most fixed effect estimates were consistent across all models for each trait (Supp. Table 4). Male nestlings were significantly heavier and had longer tarsus lengths than females; these dimorphisms increased with ontogeny. Hatch date only had a significant effect on nestling tarsus length. Larger brood size was significantly associated with lower weight for both nestlings and juveniles as well as shorter tarsus lengths at the juvenile stage. More helpers was significantly associated with increased nestling weight and tarsus lengths. More inbred nestlings had lower weight and tarsus lengths, but inbreeding did not affect juvenile traits. Larger natal territories increased nestling tarsus length and juvenile weight. Finally, age measured was not significant in any model.

## Discussion

Here we estimated indirect genetic effects from mothers, fathers, and helpers on offspring weight and tarsus length across nestling and juvenile stages. The magnitude of indirect genetic effects was negligible for most tested traits except total indirect genetic effects on nestling weight from interactions with fathers and helpers (*s*_*tot*_^*2*^ = 0.11). We predicted that fathers and helpers would contribute more to variation in offspring morphology than mothers due to their role in offspring provisioning and that indirect genetic effects would decrease across ontogeny. We discuss our findings in light of these predictions and the roles of group size and territorial resources in mediating paternal and helper genetic effects.

### Paternal and helper genetic effects contribute to variation in offspring weight

In support of our predictions, we found a significant combined effect of father and helper genetics on variation in nestling weight, with minimal maternal influence. The model with only paternal genetic effects had similar support as the model with paternal and helper effects, suggesting that individually, genetic variation in fathers contributed to variation in offspring weight, while the same is not true for helpers alone. Despite a lack of significant helper effects, the total estimated proportion of variation due to indirect genetic effects nearly doubled when helpers were added. Our results suggest that while paternal care is the primary driver of variation in nestling weight, helpers nevertheless provide a substantial synergistic effect. Our dilution parameter testing results suggest that group sizes affect the strength of indirect genetic effects such that indirect genetic effects are stronger in larger groups. We found no detectable maternal effects consistent with the observation that Florida scrub-jay mothers typically do not contribute to food provisioning of nestlings (Woolfenden and Fitzpatrick 1984). As juvenile survival is associated with nestling weight, it is possible that genetic variation in fathers and helpers may also affect variation in survival to adulthood (Mumme et al. 2015). Our results highlight the importance of considering all potential social partners when measuring indirect genetic effects.

The proportion of variation due to indirect genetic effects was not statistically different from zero for different social partners individually, but the combined effect of paternal and helper indirect genetic effects was significant and much greater than previously reported in a meta-analysis of indirect genetic effects for morphological and developmental traits (*s*_*tot*_^*2*^ = 0 - 0.02; Santostefano et al. 2025), suggesting indirect genetic effects may play a larger role in cooperative breeders. Previous estimates of indirect genetic effects in cooperatively breeding long-tailed tits (*Aegithalos caudatus*) focused on the effect of parental care on helping behavior and vice versa (Adams et al. 2015). Long-tailed tits have low indirect genetic effects on food provisioning behavior in parents from helpers, which is consistent with observations of high plasticity in food provisioning behavior and low survival between years in the studied population (Adams et al. 2015). Here, we found evidence of indirect genetic effects in a cooperative breeder but on a morphological trait at an earlier life stage. We also found evidence of stronger paternal effects on nestling weight than helper effects. Estimates of paternal effects are rare but the studies that are available suggest paternal effects are stronger in biparental systems. A study of Japanese quail did not detect paternal genetic effects on offspring mass, but that species has uniparental care from mothers (Pick et al. 2016). In burying beetles (*Nicrophorus vespilloides*) with biparental care, paternal indirect genetic effects were present and stronger than our estimates (*s*^*2*^_*g*_ = 0.20 - 0.56; Head et al. 2012). Since Florida scrub-jays perform biparental care as cooperative breeders, it is feasible that the maternal effects found in single parent systems are replaced by paternal and/or helper effects in systems with biparental or cooperative breeding systems.

### No caregiver effects on variation in tarsus length

While weight is a measure of general body size and can fluctuate over the lifetime of an individual, tarsus length is a measure of structural size and expected to be fixed following the completion of growth (Noordwijk et al. 1988). Furthermore, in altricial birds, body mass fluctuates in response to food availability, while tarsus length is relatively resistant to change (Killpack and Karasov 2012). Consistent with tarsus length being a more canalized trait, we found no significant caregiver effects on tarsus length for either developmental stage. Our results contrast with previous studies that have found maternal effects on tarsus length. Maternal effects influence tarsus length in captive Japanese quail at 2 and 4 weeks, mediated by prenatal maternal energy investment in eggs (Pick et al. 2016). Similarly, in pied flycatchers, prenatal maternal effects contribute to variation in offspring tarsus length alongside environmental factors (Potti and Merino 1994). Since we found no effect of mothers on tarsus length, it seems unlikely that prenatal maternal effects contribute to variation in tarsus lengths in our population. However, it is also possible that we have low power to detect maternal effects in tarsus length rather than a true lack of maternal effect.

### Effect of indirect genetic effects on total heritability

Given that indirect genetic effects were generally minimal for all traits except nestling weight, total heritability (*τ*^*2*^) was relatively consistent across our traits and models. Narrow-sense heritability (*h*^*2*^) was consistent across models for a given traits and developmental stage, although reduced in the presence of paternal and helper genetic effects for nestling weight. Our heritability results are consistent with our finding of significant indirect genetic effects only for nestling weight and suggest that a portion of the direct genetic variation measured in non-IGE models is attributable to contributions from social partners. In burying beetles, total heritable variation from fathers is low due to strong negative covariances between direct and indirect genetic effects (*τ*^*2*^ = -0.36 - 0.12; Head et al. 2012). A recent meta-analysis found total heritable variation tends to be 66% greater than estimates of narrow-sense heritability, implying that social partners often increase the amount of genetic variation available for natural selection to act on (Santostefano et al. 2025). Here, we found that total heritability was greater than narrow-sense heritability for nestling weight, implying that some genetic variation in nestling weight is misattributed to additive genetic effects when paternal and helper effects are not estimated (Kruuk 2004). Our results suggest a potentially strong role of fathers and helpers and not mothers or the nestling itself on evolutionary trajectories of offspring weight.

### Relative importance of direct and indirect genetic effects across ontogeny

Direct genetic effects increased across ontogeny regardless of the presence of indirect genetic effects. We found juvenile traits were similarly heritable to a meta-analysis of morphological traits (average *h*^*2*^ = 0.46; Mousseau and Roff 1987), whereas nestling traits were substantially less heritable. Our findings align with results reported in *Ovis* and *Bos* species (Wilson and Réale 2006), red deer (Gauzere et al. 2020), and Japanese Quail (Vedder et al. 2023). Consistent with previous studies that have shown a concurrent decrease in indirect genetic effects with ontogeny (Vedder et al. 2023), we found a decrease in paternal and helper effects on weight from nestling to juvenile stages. We expected to find lower indirect genetic effects for juveniles at nutritional independence because at that age, offspring are no longer reliant on their parents and helpers for food (Woolfenden and Fitzpatrick 1984). However, it is also possible that our smaller sample sizes for juvenile traits compared to nestling traits, paired with lower variances in juvenile traits, reduced our capacity to detect indirect genetic effects at the later life stage. Estimates of heritability and indirect genetic effects across ontogeny remain rare and further studies are needed to better understand the changing role of caregivers as offspring develop.

### Caveats and Future Directions

Estimates of indirect genetic effects in the wild are difficult to obtain in part due to the large sample sizes required to robustly estimate quantitative genetic parameters (Charmantier et al. 2014; Baud et al. 2022; Santostefano et al. 2025). While our dataset was large compared to many other studies of wild populations, we may still have had insufficient power, particularly for juvenile traits (Table 1). We note that we only detected significant indirect genetic effects for nestling weight, the trait with the largest sample size in our study. Our low power to estimate indirect genetic effects also prevented us from estimating covariances between direct and indirect genetic effects in every trait for every social partner, particularly in models in which indirect genetic variance was near 0. For those models, we were unable to completely measure total heritable variation (Bijma 2011) or infer whether indirect genetic effects constrain or amplify trait evolution. Increased sample sizes would improve our estimates of variances and covariances. Additionally, we did not measure the effect of spatial autocorrelation that would occur due to limited dispersal and natal philopatry which are known to upwardly bias estimates of narrow-sense heritability in red deer (Stopher et al. 2012) and may bias indirect environmental effects (Santostefano et al. 2021). Future studies should incorporate spatial autocorrelation to assess these effects.

Here we relied on variance partitioning models and the pedigree to infer indirect genetic effects on offspring traits. These models do not provide information on which specific traits mediate indirect genetic effects unlike trait-based models, which require more extensive data (Baud et al. 2022). Recently, we generated a high quality Florida scrub-jay reference genome (Romero et al. 2024), and we are now genotyping nearly 4,000 individuals, which will allow us to leverage polygenic risk score approaches in the future to test the specific effect of traits such as food provisioning behavior. Polygenic risk score approaches are both trait-based and genetic, where genotypes and effect sizes from genome-wide association studies are used to predict phenotypes (Baud et al. 2022). For example, Kong et al. (2018) compared regressions of offspring phenotypes on polygenic risk scores calculated from transmitted and non-transmitted alleles to show that indirect genetic effects exist for educational attainment in humans.Generally, the sample sizes required to estimate direct and indirect genetic effects for each single nucleotide polymorphism are very large, but recent statistical developments offer increased power (Wu et al. 2021)

### Summary

We found that in Florida scrub-jays, paternal and helper genotypes affect variation in nestling weight. Helper genetic effects are likely mediated by the number of helpers present in the population and are secondary to the effect of fathers. We found no evidence of maternal effects on offspring traits in Florida scrub-jays, consistent with known patterns of food provisioning behavior, and no indirect effects on tarsus length in any of our models. We also provide evidence for increasing additive genetic variation across ontogeny. Our study provides new insights into the role of helpers in cooperatively breeding systems in trait evolution and emphasizes the importance of considering multiple social partners and interactions among them when modeling evolution in cooperatively-breeding species. While studies of indirect genetic effects in wild populations remain rare due to data limitations, emerging genomic and statistical approaches may facilitate more robust estimates of indirect genetic effects across diverse study systems, ultimately enhancing our understanding of offspring trait evolution.

## Funding

This work was supported by NIH grant 1R35GM133412 to NC and a Schwartz Discover Grant to GS.

## Conflict of Interest Statement

GS, DT, EC, TDB, SB, RB, JWF and NC declare that they have no conflicts of interest.

## Acknowledgements

Thank you to all the interns and staff at Archbold Biological Station who collected demographic data on the Florida scrub-jay, as well as to the members of the Chen Lab and the Avian Ecology Program at Archbold Biological Station for their valuable feedback.

Model 1 is a null model with no random effects. In model 2, natal year was fitted. In model 3, natal year and nest were fitted. In model 4, additive genetic effects were added. In model 5, social partner identity was added as an environmental effect for each social partner separately and in model 6, social partner genetic effects were added. Model 7 included a covariance structure between direct and indirect genetic effects. In model M+D, maternal and paternal effects were included. In model M+H, maternal and paternal effects were included. In model D+H, paternal and helper effects were included. In model M+D+H, all three social partners were included in a single model.

## Notes

### Competing Interest Statement

The authors have declared no competing interest.

## References

Adams, M. J., M. R. Robinson, M.-E. Mannarelli, and B. J. Hatchwell. 2015. Social genetic and social environment effects on parental and helper care in a cooperatively breeding bird. Proc Biol Sci 282:20150689.

Baud, A., S. McPeek, N. Chen, and K. A. Hughes. 2022. Indirect Genetic Effects: A Cross-disciplinary Perspective on Empirical Studies. J Hered 113:1–15.

Bijma, P. 2011. A general definition of the heritable variation that determines the potential of a population to respond to selection. Genetics 189:1347–1359.

Bijma, P. 2010. Estimating Indirect Genetic Effects: Precision of Estimates and Optimum Designs. Genetics 186:1013–1028.

Bijma, P. 2014. The quantitative genetics of indirect genetic effects: a selective review of modelling issues. Heredity (Edinb) 112:61–69.

Bijma, P., and M. J. Wade. 2008. The joint effects of kin, multilevel selection and indirect genetic effects on response to genetic selection. Journal of Evolutionary Biology 21:1175–1188.

Bleakley, B. H., and E. D. III Brodie. 2009. Indirect genetic effects influence antipredator behavior in guppies: estimates of the coefficient of interaction psi and the inheritance of reciprocity. Evol 63:1796–1806.

Butler, D. G., B. R. Cullis, A. R. Gilmour, B. G. Gogel, and R. Thompson. 2023. ASReml-R Reference Manual Version 4.2. VSN International Ltd., Hemel Hempstead, HP2 4TP, UK.

Charmantier, E. by A., D. Garant, and and L. E. B. Kruuk (eds). 2014. Quantitative Genetics in the Wild. Oxford University Press, Oxford, New York.

Chen, N., E. J. Cosgrove, R. Bowman, J. W. Fitzpatrick, and A. G. Clark. 2016. Genomic Consequences of Population Decline in the Endangered Florida scrub-jay. Current Biology 26:2974–2979.

Cheverud, J. M., and A. J. Moore. 1994. Quantitative genetic and the role of the environment provided by relatives in behavioral evolution. Pp. 67–100 in Quantitative Genetic Studies of Behavioral Evolution. University of Chicago Press.

Coster, A. 2022. pedigree: Pedigree Functions.

Crean, A. J., and R. Bonduriansky. 2014. What is a paternal effect? Trends in Ecology & Evolution 29:554–559. Elsevier.

Falconer, D. S., and T. F. MacKay. 1996. Introduction to Quantitative Genetics. Longman Publishing Group, Harlow.

Fisher, D. N., A. J. Wilson, S. Boutin, B. Dantzer, J. E. Lane, D. W. Coltman, J. C. Gorrell, and A. G. McAdam. 2019. Social effects of territorial neighbours on the timing of spring breeding in North American red squirrels. J Evol Biol 32:559–571.

Gauzere, J., J. M. Pemberton, S. Morris, A. Morris, L. E. B. Kruuk, and C. A. Walling. 2020. The genetic architecture of maternal effects across ontogeny in the red deer. Evol 74:1378–1391.

Hadfield, J. D. 2010. MCMC Methods for Multi-Response Generalized Linear Mixed Models: The MCMCglmm R Package. Journal of Statistical Software 33:1–22.

Head, M. L., L. K. Berry, N. J. Royle, and A. J. Moore. 2012. Paternal care: direct and indirect genetic effects of fathers on offspring performance. Evolution 66:3570–3581.

Hunt, J., and L. W. Simmons. 2000. Maternal and paternal effects on offspring phenotyped in the dung beetle Onthophagus taurus. Evol 54:936–941.

Killpack, T. L., and W. H. Karasov. 2012. Growth and development of house sparrows (Passer domesticus) in response to chronic food restriction throughout the nestling period. J Exp Biol 215:1806–1815.

Kong, A., G. Thorleifsson, M. L. Frigge, B. J. Vilhjalmsson, A. I. Young, T. E. Thorgeirsson, S. Benonisdottir, A. Oddsson, B. V. Halldorsson, G. Masson, D. F. Gudbjartsson, A. Helgason, G. Bjornsdottir, U. Thorsteinsdottir, and K. Stefansson. 2018. The nature of nurture: Effects of parental genotypes. Science 359:424–428. American Association for the Advancement of Science.

Kruuk, L. E. B. 2004. Estimating genetic parameters in natural populations using the “animal model”. Philos Trans R Soc Lond B Biol Sci 359:873–890.

McAdam, A. G., S. Boutin, D. Réale, and D. Berteaux. 2002. Maternal effects and the potential for evolution in a natural population of animals. Evol 56:846–851.

McAdam, A. G., D. Garant, and A. J. Wilson. 2014. The effects of others’ genes: maternal and other indirect genetic effects. P. 0 in A. Charmantier, D. Garant, and L. E. B. Kruuk, eds. Quantitative Genetics in the Wild. Oxford University Press.

McGlothlin, J. W., and E. D. Brodie III. 2009. How to Measure Indirect Genetic Effects: The Congruence of Trait-Based and Variance-Partitioning Approaches. Evolution 63:1785–1795.

Moore, A. J., E. D. III Brodie, and J. B. Wolf. 1997. Interacting phenotypes and the evolutionary process: I. Direct and indirect genetic effects of social interactions. Evol 51:1352–1362.

Mousseau, T. A., and D. A. Roff. 1987. Natural selection and the heritability of fitness components. Heredity (Edinb) 59 (Pt 2):181–197.

Mumme, R. L., R. Bowman, M. S. Pruett, and J. W. Fitzpatrick. 2015. Natal territory size, group size, and body mass affect lifetime fitness in the cooperatively breeding Florida scrub-jay. Auk 132:634–646.

Noordwijk, A. J. V., J. H. V. Balen, and W. Scharloo. 1988. Heritability of body size in a natural population of the Great Tit (Parus major) and its relation to age and environmental conditions during growth. Genetics Research 51:149–162.

Pick, J. L., C. Ebneter, P. Hutter, and B. Tschirren. 2016. Disentangling Genetic and Prenatal Maternal Effects on Offspring Size and Survival. The American Naturalist 188:628–639. The University of Chicago Press.

Potti, J., and S. Merino. 1994. Heritability estimates and maternal effects on tarsus length in pied flycatchers, Ficedula hypoleuca. Oecologia 100:331–338.

Quinn, J. S., G. E. Woolfenden, J. W. Fitzpatrick, and B. N. White. 1999. Multi-locus DNA fingerprinting supports genetic monogamy in Florida scrub-jays. Behav Ecol Sociobiol 45:1–10.

R Core Team. 2023. R: A Language and Environment for Statistical Computing. Vienna, Austria.

Romero, F. G., F. E. G. Beaudry, E. Hovmand Warner, T. N. Nguyen, J. W. Fitzpatrick, and N. Chen. 2024. A new high-quality genome assembly and annotation for the threatened Florida scrub-jay (Aphelocoma coerulescens). G3 Genes|Genomes|Genetics, doi: 10.1093/g3journal/jkae232.

Santostefano, F., H. Allegue, D. Garant, P. Bergeron, and D. Réale. 2021. Indirect genetic and environmental effects on behaviors, morphology, and life-history traits in a wild Eastern chipmunk population. Evol 75:1492–1512.

Santostefano, F., M. Moiron, A. Sánchez-Tójar, and D. N. Fisher. 2025. Indirect genetic effects increase the heritable variation available to selection and are largest for behaviors: a meta-analysis. Evol Lett 9:89–104.

Self, S. G., and K.-Y. Liang. 1987. Asymptotic Properties of Maximum Likelihood Estimators and Likelihood Ratio Tests Under Nonstandard Conditions. Journal of the American Statistical Association 82:605–610. [American Statistical Association, Taylor & Francis, Ltd.].

Sinnwell, J., T. Therneau, D. Schaid, E. Atkinson, and C. Mester. 2024. kinship2: Pedigree Functions.

Skutch, A. F. 1935. Helpers at the Nest. Auk 52:257–273.

Stopher, K. V., D. H. Nussey, T. H. Clutton-Brock, F. Guinness, A. Morris, and J. M. Pemberton. 2012. Re-mating across years and intralineage polygyny are associated with greater than expected levels of inbreeding in wild red deer. J Evol Biol 25:2457–2469.

Suh, Y. H., R. Bowman, and J. W. Fitzpatrick. 2022. Staging to join non-kin groups in a classical cooperative breeder, the Florida scrub-jay. Journal of Animal Ecology 91:970–982.

Townsend, A. K., R. Bowman, J. W. Fitzpatrick, M. Dent, and I. J. Lovette. 2011. Genetic monogamy across variable demographic landscapes in cooperatively breeding Florida scrub-jays. Behav Ecol 22:464–470.

Vedder, O., B. Tschirren, E. Postma, and M. Moiron. 2023. Rapid decline of prenatal maternal effects with age is independent of postnatal environment in a precocial bird. Evol 77:2484–2491.

White, S. J., and A. J. Wilson. 2019. Evolutionary genetics of personality in the Trinidadian guppy I: maternal and additive genetic effects across ontogeny. Heredity 122:1–14. Nature Publishing Group.

Wilson, A. J., and D. Réale. 2006. Ontogeny of Additive and Maternal Genetic Effects: Lessons from Domestic Mammals. The American Naturalist 167:E23–E38. The University of Chicago Press.

Wilson, A. J., D. Réale, M. N. Clements, M. M. Morrissey, E. Postma, C. A. Walling, L. E. B. Kruuk, and D. H. Nussey. 2010. An ecologist’s guide to the animal model. Journal of Animal Ecology 79:13–26.

Wolf, J. B., E. D. Brodie III, J. M. Cheverud, A. J. Moore, and M. J. Wade. 1998. Evolutionary consequences of indirect genetic effects. Trends in Ecology & Evolution 13:64–69.

Woolfenden, G. E. 1975. Florida Scrub Jay Helpers at the Nest. Auk 92:1–15.

Woolfenden, G. E., and J. W. Fitzpatrick. 1984. The Florida Scrub Jay: Demography of a Cooperative-Breeding Bird. Princeton University Press.

Wu, Y., X. Zhong, Y. Lin, Z. Zhao, J. Chen, B. Zheng, J. J. Li, J. M. Fletcher, and Q. Lu. 2021. Estimating genetic nurture with summary statistics of multigenerational genome-wide association studies. Proceedings of the National Academy of Sciences 118:e2023184118. Proceedings of the National Academy of Sciences.

