## Supplemental Files for "Indirect genetic effects across ontogeny in an avian cooperative breeder"

### Supplementary Figures and Tables

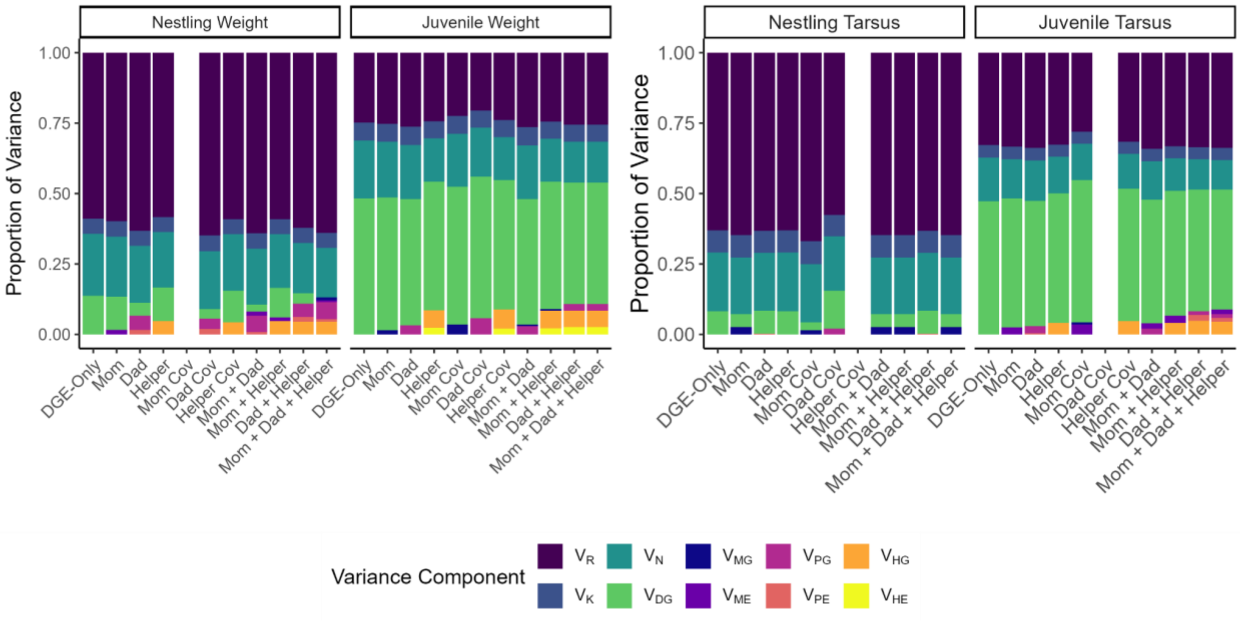

**Supplementary Figure 1:** Estimated variance components as proportions of total phenotypic variance in offspring weight and tarsus length at the nestling and juvenile stages. For each trait, we ran DGE-only, maternal, paternal, and helper models, as well as models with multiple social partners to test the effect of different social partners. Models in which an unstructured covariance was fitted between additive genetic variance and social partner genetic variance are labeled with "Cov". Three models with covariance terms did not converge and are therefore omitted from the plot. Colors represent different variance components. $V_{DG}$ is the direct genetic effect, $V_{MG}$ is the maternal genetic effect, $V_{ME}$ is the maternal environmental effect, $V_{PG}$ is the paternal genetic effect, $V_{PE}$ is the paternal environmental effect, $V_{HG}$ is the helper genetic effect, $V_{HE}$ is the helper environmental effect, $V_N$ is the natal nest effect, $V_K$ is the natal year effect, and $V_R$ is the residual. For full model results, see Table 2.

**Supplementary Table 1. Tabular Form of Narrow-Sense Heritability, Relative Total Heritable Variation, and Social Effect Parameters.** Standard errors are in parentheses when available. Significant values are in bold face determined by 2*standard error is greater than 0. $h^2$ is narrow-sense heritability. $\tau^2$ is proportional total heritable variance. TBV is total heritable variance. $s_g^2$ is the scaled parameter for indirect genetic effects calculated as $V_{SG}/V_P$. $s_e^2$ is the scaled parameter for indirect environmental effects calculated as $V_{SE}/V_P$. $s_{tot}^2$ is the scaled parameter for total indirect effects calculated as $(V_{SG} + V_{SE})/V_P$. M, D, and H refer to the family members included in each model (mom, dad, helper).

| Trait | Model | $h^2$ | $\tau^2$ | $s_g^2$ | $s_e^2$ | $s_{tot}^2$ |
|---|---|---|---|---|---|---|
| Nestling Weight | DGE Only | **0.14 (0.05)** | | | | |
| | M | **0.12 (0.04)** | **0.12 (0.04)** | <0.0001 | 0.016 (0.019) | 0.016 (0.019) |
| | D | 0.046 (0.042) | **0.097 (0.039)** | 0.05 (0.033) | 0.015 (0.027) | **0.066 (0.024)** |
| | H | **0.12 (0.034)** | **0.15 (0.035)** | 0.047 (0.025) | <0.0001 | 0.047 (0.025) |
| | D Cov | 0.03 (0.05) | **0.11 (0.04)** | 0.02 (0.04) | 0.04 (0.05) | 0.05 (0.03) |
| | H Cov | **0.11 (0.04)** | **0.11 (0.04)** | 0.04 (0.03) | <0.0001 | 0.04 (0.03) |
| | M+D | 0.025 (0.049) | 0.083 (0.042) | 0.058 (0.034) | 0.022 (0.029) | **0.08 (0.032)** |
| | M+H | **0.1 (0.039)** | **0.14 (0.04)** | 0.046 (0.025) | 0.014 (0.018) | **0.061 (0.03)** |
| | D+H | 0.036 (0.041) | **0.11 (0.04)** | **0.093 (0.04)** | 0.062 (0.036) | **0.11 (0.033)** |
| | M+D+H | <0.0001 | **0.1 (0.04)** | **0.12 (0.043)** | 0.014 (0.036) | **0.13 (0.03)** |
| Juvenile Weight | DGE Only | **0.48 (0.06)** | | | | |
| | M | **0.47 (0.073)** | **0.49 (0.066)** | 0.014 (0.032) | <0.0001 | 0.014 (0.032) |
| | D | **0.45 (0.078)** | **0.48 (0.067)** | 0.032 (0.037) | <0.0001 | 0.032 (0.037) |
| | H | **0.46 (0.066)** | **0.51 (0.078)** | 0.063 (0.052) | 0.023 (0.072) | 0.086 (0.054) |
| | M Cov | **0.49 (0.08)** | **0.48 (0.1)** | -0.12 (0.19) | 0.09 (0.17) | -0.03 (0.15) |
| | D Cov | **0.5 (0.09)** | **0.5 (0.07)** | -0.19 (0.23) | 0.15 (0.14) | -0.04 (0.15) |
| | H Cov | **0.46 (0.07)** | **0.46 (0.07)** | 0.07 (0.06) | 0.02 (0.07) | 0.09 (0.06) |
| | M+D | **0.45 (0.083)** | **0.48 (0.068)** | 0.035 (0.044) | <0.0001 | 0.035 (0.044) |
| | M+H | **0.45 (0.073)** | **0.52 (0.078)** | 0.068 (0.059) | 0.062 (0.052) | 0.09 (0.061) |
| | D+H | **0.43 (0.077)** | **0.51 (0.079)** | 0.081 (0.06) | 0.058 (0.051) | 0.11 (0.063) |
| | M+D+H | **0.43 (0.081)** | **0.51 (0.079)** | 0.081 (0.063) | 0.027 (0.071) | 0.11 (0.066) |
| Nestling Tarsus | DGE Only | **0.08 (0.01)** | | | | |
| | M | 0.046 (0.044) | **0.072 (0.034)** | 0.026 (0.024) | <0.0001 | 0.026 (0.024) |
| | D | **0.081 (0.04)** | **0.081 (0.04)** | <0.0001 | 0.0028 (0.02) | 0.0028 (0.02) |

| | | | | | | |
|---|---|---|---|---|---|---|
| | H | **0.083 (0.036)** | **0.083 (0.036)** | <0.0001 | <0.0001 | <0.0001 |
| | M Cov | 0.02 (0.06) | 0.08 (0.04) | 0.02 (0.05) | 0.02 (0.04) | 0.03 (0.03) |
| | D Cov | 0.13 (0.07) | 0.07 (0.04) | -0.06 (0.06) | 0.03 (0.05) | -0.03 (0.04) |
| | M+D | 0.046 (0.044) | **0.072 (0.034)** | 0.026 (0.024) | <0.0001 | 0.026 (0.024) |
| | M+H | 0.046 (0.044) | **0.072 (0.034)** | 0.026 (0.024) | <0.0001 | 0.026 (0.035) |
| | D+H | **0.081 (0.04)** | **0.081 (0.04)** | <0.0001 | 0.0028 (0.02) | 0.0028 (0.034) |
| | M+D+H | **0.046 (0.044)** | **0.072 (0.034)** | 0.026 (0.024) | <0.0001 | 0.026 (0.024) |
| Juvenile Tarsus | DGE Only | **0.47 (0)** | | | | |
| | M | **0.46 (0.068)** | **0.46 (0.068)** | <0.0001 | 0.024 (0.03) | 0.024 (0.03) |
| | D | **0.45 (0.076)** | **0.47 (0.071)** | 0.026 (0.056) | 0.0039 (0.044) | 0.03 (0.039) |
| | H | **0.46 (0.063)** | **0.5 (0.064)** | 0.041 (0.035) | <0.0001 | 0.041 (0.035) |
| | M Cov | **0.5 (0.08)** | **0.43 (0.08)** | -0.16 (0.23) | 0.03 (0.19) | -0.14 (0.17) |
| | H Cov | **0.47 (0.07)** | **0.47 (0.07)** | 0.05 (0.04) | <0.0001 | 0.05 (0.04) |
| | M+D | **0.44 (0.08)** | **0.46 (0.073)** | 0.021 (0.056) | 0.02 (0.048) | 0.04 (0.043) |
| | M+H | **0.44 (0.068)** | **0.48 (0.07)** | 0.042 (0.035) | 0.066 (0.046) | 0.066 (0.046) |
| | D+H | **0.43 (0.077)** | **0.49 (0.072)** | 0.061 (0.065) | 0.069 (0.062) | 0.082 (0.054) |
| | M+D+H | **0.43 (0.08)** | **0.48 (0.074)** | 0.06 (0.065) | 0.029 (0.049) | 0.088 (0.056) |

**Supplementary Table 2. Dilution parameter testing results without relatedness.** Results are ordered by smallest to largest AIC for each model type. ΔAIC is measured from the best performing model. Differences between model fit for tarsus length models were too small (<0.0001) to show here.

| Trait | Social Partner | Model | LogL | AIC | ΔAIC | LogL | AIC | ΔAIC |
|---|---|---|---|---|---|---|---|---|
| | | | Nestling | | | Juvenile | | |
| Weight | Helper | d1 | -7619 | 15251 | 0 | -2392 | 4796.44 | 0.01 |
| | | d0.9 | -7620 | 15251.37 | 0.37 | -2392 | 4796.44 | 0 |
| | | d0.8 | -7620 | 15251.76 | 0.77 | -2392 | 4796.44 | 0.01 |
| | | d0.7 | -7620 | 15252.17 | 1.17 | -2392 | 4796.47 | 0.03 |
| | | d0.6 | -7620 | 15252.58 | 1.58 | -2392 | 4796.51 | 0.07 |
| | | d0.5 | -7620 | 15252.98 | 1.98 | -2392 | 4796.56 | 0.12 |
| | | d0.4 | -7621 | 15253.38 | 2.38 | -2392 | 4796.63 | 0.19 |
| | | d0.3 | -7621 | 15253.77 | 2.77 | -2392 | 4796.71 | 0.27 |
| | | d0.2 | -7621 | 15254.14 | 3.14 | -2392 | 4796.8 | 0.37 |
| | | d0.1 | -7621 | 15254.48 | 3.48 | -2392 | 4796.91 | 0.47 |
| | | d0 | -7621 | 15254.8 | 3.8 | -2393 | 4797.03 | 0.59 |
| Tarsus | | d0 | -2762 | 5536.22 | 0 | -652 | 1315.61 | 0 |
| | | d0.1 | -2762 | 5536.22 | 0 | -652 | 1315.74 | 0.13 |
| | | d0.2 | -2762 | 5536.22 | 0 | -652 | 1315.79 | 0.17 |
| | | d0.3 | -2762 | 5536.22 | 0 | -652 | 1315.81 | 0.2 |
| | | d0.4 | -2762 | 5536.22 | 0 | -652 | 1315.84 | 0.23 |
| | | d0.5 | -2762 | 5536.22 | 0 | -652 | 1315.87 | 0.26 |
| | | d0.6 | -2762 | 5536.22 | 0 | -652 | 1315.88 | 0.27 |
| | | d0.7 | -2762 | 5536.22 | 0 | -652 | 1315.93 | 0.31 |
| | | d0.8 | -2762 | 5536.22 | 0 | -652 | 1315.98 | 0.37 |
| | | d0.9 | -2762 | 5536.22 | 0 | -652 | 1316 | 0.39 |
| | | d1 | -2762 | 5536.22 | 0 | -652 | 1316.04 | 0.43 |
| Weight | Dad + Helper | d1 | -7616 | 15247.12 | 0 | -2392 | 4799.93 | 0 |
| | | d0.9 | -7616 | 15247.41 | 0.29 | -2392 | 4799.93 | 0 |
| | | d0.8 | -7616 | 15247.72 | 0.6 | -2392 | 4799.94 | 0.02 |
| | | d0.7 | -7616 | 15248.04 | 0.92 | -2392 | 4799.97 | 0.04 |
| | | d0.6 | -7616 | 15248.37 | 1.25 | -2392 | 4800.02 | 0.09 |
| | | d0.5 | -7616 | 15248.71 | 1.59 | -2392 | 4800.07 | 0.15 |
| | | d0.4 | -7617 | 15249.04 | 1.92 | -2392 | 4800.14 | 0.22 |
| | | d0.3 | -7617 | 15249.37 | 2.25 | -2392 | 4800.23 | 0.3 |
| | | d0.2 | -7617 | 15249.68 | 2.57 | -2392 | 4800.33 | 0.4 |
| | | d0.1 | -7617 | 15249.99 | 2.87 | -2392 | 4800.43 | 0.51 |
| | | d0 | -7617 | 15250.27 | 3.15 | -2394 | 4803.22 | 3.3 |
| Tarsus | | d0 | -2762 | 5540.2 | 0 | -651 | 1318.67 | 0 |
| | | d0.1 | -2762 | 5540.2 | 0 | -651 | 1318.83 | 0.16 |

| | | | | | | | | |
|---|---|---|---|---|---|---|---|---|
| | | d0.2 | -2762 | 5540.2 | 0 | -651 | 1318.98 | 0.31 |
| | | d0.3 | -2762 | 5540.2 | 0 | -651 | 1318.99 | 0.32 |
| | | d0.4 | -2762 | 5540.2 | 0 | -652 | 1319.02 | 0.34 |
| | | d0.5 | -2762 | 5540.2 | 0 | -652 | 1319.06 | 0.39 |
| | | d0.6 | -2762 | 5540.2 | 0 | -652 | 1319.12 | 0.44 |
| | | d0.7 | -2762 | 5540.2 | 0 | -652 | 1319.16 | 0.48 |
| | | d0.8 | -2762 | 5540.2 | 0 | -652 | 1319.18 | 0.51 |
| | | d0.9 | -2762 | 5540.2 | 0 | -652 | 1319.25 | 0.57 |
| | | d1 | -2762 | 5540.2 | 0 | -652 | 1319.32 | 0.64 |
| Weight | Mom + Helper | d1 | -7619 | 15254.35 | 0 | -2392 | 4800.4 | 0 |
| | | d0.9 | -7619 | 15254.7 | 0.35 | -2392 | 4800.41 | 0.01 |
| | | d0.8 | -7620 | 15255.07 | 0.72 | -2392 | 4800.41 | 0 |
| | | d0.7 | -7620 | 15255.44 | 1.09 | -2392 | 4800.44 | 0.03 |
| | | d0.6 | -7620 | 15255.83 | 1.48 | -2392 | 4800.47 | 0.07 |
| | | d0.5 | -7620 | 15256.21 | 1.86 | -2392 | 4800.53 | 0.13 |
| | | d0.4 | -7620 | 15256.58 | 2.23 | -2392 | 4800.59 | 0.19 |
| | | d0.3 | -7620 | 15256.95 | 2.6 | -2392 | 4800.67 | 0.27 |
| | | d0.2 | -7621 | 15257.29 | 2.94 | -2392 | 4800.77 | 0.37 |
| | | d0.1 | -7621 | 15257.62 | 3.27 | -2392 | 4800.87 | 0.47 |
| | | d0 | -7621 | 15257.93 | 3.58 | -2392 | 4800.98 | 0.58 |
| Tarsus | | d0 | -2761 | 5538.81 | 0 | -651 | 1318.83 | 0 |
| | | d0.1 | -2761 | 5538.81 | 0 | -651 | 1318.89 | 0.06 |
| | | d0.2 | -2761 | 5538.81 | 0 | -651 | 1318.92 | 0.09 |
| | | d0.3 | -2761 | 5538.81 | 0 | -651 | 1318.96 | 0.14 |
| | | d0.4 | -2761 | 5538.81 | 0 | -651 | 1318.96 | 0.13 |
| | | d0.5 | -2761 | 5538.81 | 0 | -652 | 1319.01 | 0.18 |
| | | d0.6 | -2761 | 5538.81 | 0 | -652 | 1319.07 | 0.24 |
| | | d0.7 | -2761 | 5538.81 | 0 | -652 | 1319.1 | 0.27 |
| | | d0.8 | -2761 | 5538.81 | 0 | -652 | 1319.13 | 0.3 |
| | | d0.9 | -2761 | 5538.81 | 0 | -652 | 1319.2 | 0.37 |
| | | d1 | -2761 | 5538.81 | 0 | -652 | 1319.24 | 0.41 |
| Weight | Mom + Dad + Helper | d1 | -7615 | 15250.84 | 0 | -2392 | 4803.93 | 0 |
| | | d0.9 | -7616 | 15251.13 | 0.28 | -2392 | 4803.93 | 0 |
| | | d0.8 | -7616 | 15251.43 | 0.59 | -2392 | 4803.94 | 0.02 |
| | | d0.7 | -7616 | 15251.75 | 0.9 | -2392 | 4803.97 | 0.04 |
| | | d0.6 | -7616 | 15252.07 | 1.23 | -2392 | 4804.02 | 0.09 |
| | | d0.5 | -7616 | 15252.4 | 1.55 | -2392 | 4804.07 | 0.15 |
| | | d0.4 | -7616 | 15252.72 | 1.88 | -2392 | 4804.14 | 0.22 |
| | | d0.3 | -7617 | 15253.04 | 2.2 | -2392 | 4804.23 | 0.3 |
| | | d0.2 | -7617 | 15253.36 | 2.51 | -2392 | 4804.33 | 0.4 |
| | | d0.1 | -7617 | 15253.65 | 2.81 | -2392 | 4804.43 | 0.51 |
| | | d0 | -7617 | 15253.93 | 3.09 | -2394 | 4807.2 | 3.28 |
| Tarsus | | d0 | -2761 | 5542.81 | 0 | -651 | 1322.41 | 0 |

| | | | | | | | | |
|---|---|---|---|---|---|---|---|---|
| | | d0.1 | -2761 | 5542.81 | 0 | -651 | 1322.56 | 0.15 |
| | | d0.2 | -2761 | 5542.81 | 0 | -651 | 1322.58 | 0.17 |
| | | d0.3 | -2761 | 5542.81 | 0 | -651 | 1322.62 | 0.2 |
| | | d0.4 | -2761 | 5542.81 | 0 | -651 | 1322.66 | 0.25 |
| | | d0.5 | -2761 | 5542.81 | 0 | -651 | 1322.72 | 0.31 |
| | | d0.6 | -2761 | 5542.81 | 0 | -651 | 1322.72 | 0.3 |
| | | d0.7 | -2761 | 5542.81 | 0 | -651 | 1322.78 | 0.37 |
| | | d0.8 | -2761 | 5542.81 | 0 | -651 | 1322.85 | 0.43 |
| | | d0.9 | -2761 | 5542.81 | 0 | -651 | 1322.87 | 0.46 |
| | | d1 | -2761 | 5542.81 | 0 | -651 | 1322.92 | 0.51 |

**Supplementary Table 3: Dilution parameter testing results with relatedness.** Results are ordered by smallest to largest AIC for each model type. ΔAIC is measured from the best performing model. Differences between model fit for tarsus length models were too small (<0.0001) to show here.

| Trait | Social Partner | Model | LogL | AIC | ΔAIC | LogL | AIC | ΔAIC |
|---|---|---|---|---|---|---|---|---|
| | | | | Nestling | | | Juvenile | |
| Weight | Helper | d1_rel | -7621 | 15254.99 | 3.99 | -2394 | 4799.88 | 3.45 |
| | | d0.9_rel | -7622 | 15255.15 | 4.15 | -2394 | 4799.89 | 3.45 |
| | | d0.8_rel | -7622 | 15255.31 | 4.31 | -2394 | 4799.89 | 3.46 |
| | | d0.7_rel | -7622 | 15255.47 | 4.47 | -2394 | 4799.9 | 3.46 |
| | | d0.6_rel | -7622 | 15255.62 | 4.63 | -2394 | 4799.91 | 3.48 |
| | | d0.5_rel | -7622 | 15255.77 | 4.77 | -2394 | 4799.93 | 3.49 |
| | | d0.4_rel | -7622 | 15255.9 | 4.9 | -2394 | 4799.95 | 3.51 |
| | | d0.3_rel | -7622 | 15256.02 | 5.03 | -2394 | 4799.97 | 3.54 |
| | | d0.2_rel | -7622 | 15256.13 | 5.13 | -2394 | 4800 | 3.57 |
| | | d0.1_rel | -7622 | 15256.22 | 5.22 | -2394 | 4800.04 | 3.6 |
| | | d0_rel | -7622 | 15256.29 | 5.29 | -2394 | 4800.07 | 3.64 |
| Tarsus | | d0_rel | -2762 | 5536.22 | 0 | -652 | 1316.1 | 0.49 |
| | | d0.1_rel | -2762 | 5536.22 | 0 | -652 | 1316.14 | 0.52 |
| | | d0.2_rel | -2762 | 5536.22 | 0 | -652 | 1316.17 | 0.55 |
| | | d0.3_rel | -2762 | 5536.22 | 0 | -652 | 1316.24 | 0.62 |
| | | d0.4_rel | -2762 | 5536.22 | 0 | -652 | 1316.27 | 0.66 |
| | | d0.5_rel | -2762 | 5536.22 | 0 | -652 | 1316.31 | 0.7 |
| | | d0.6_rel | -2762 | 5536.22 | 0 | -652 | 1316.4 | 0.78 |
| | | d0.7_rel | -2762 | 5536.22 | 0 | -652 | 1316.52 | 0.91 |
| | | d0.8_rel | -2762 | 5536.22 | 0 | -652 | 1316.64 | 1.02 |
| | | d0.9_rel | -2762 | 5536.22 | 0 | -652 | 1316.75 | 1.14 |
| | | d1_rel | -2762 | 5536.22 | 0 | -652 | 1316.85 | 1.24 |
| Weight | Dad + Helper | d1_rel | -7617 | 15250.18 | 3.06 | -2392 | 4800.55 | 0.62 |
| | | d0.9_rel | -7617 | 15250.34 | 3.22 | -2394 | 4803.22 | 3.29 |
| | | d0.8_rel | -7617 | 15250.5 | 3.38 | -2394 | 4803.22 | 3.29 |
| | | d0.7_rel | -7617 | 15250.67 | 3.55 | -2394 | 4803.22 | 3.3 |
| | | d0.6_rel | -7617 | 15250.84 | 3.72 | -2394 | 4803.23 | 3.31 |
| | | d0.5_rel | -7618 | 15251.01 | 3.89 | -2394 | 4803.24 | 3.32 |
| | | d0.4_rel | -7618 | 15251.16 | 4.05 | -2394 | 4803.26 | 3.33 |
| | | d0.3_rel | -7618 | 15251.31 | 4.2 | -2394 | 4803.28 | 3.35 |
| | | d0.2_rel | -7618 | 15251.45 | 4.33 | -2394 | 4803.3 | 3.38 |
| | | d0.1_rel | -7618 | 15251.58 | 4.46 | -2394 | 4803.33 | 3.4 |
| | | d0_rel | -7618 | 15251.69 | 4.57 | -2394 | 4803.36 | 3.43 |
| Tarsus | | d0_rel | -2762 | 5540.2 | 0 | -652 | 1319.32 | 0.65 |
| | | d0.1_rel | -2762 | 5540.2 | 0 | -652 | 1319.4 | 0.73 |
| | | d0.2_rel | -2762 | 5540.2 | 0 | -652 | 1319.48 | 0.8 |

| | | | | | | | | |
|---|---|---|---|---|---|---|---|---|
| | | d0.3_rel | -2762 | 5540.2 | 0 | -652 | 1319.48 | 0.81 |
| | | d0.4_rel | -2762 | 5540.2 | 0 | -652 | 1319.57 | 0.89 |
| | | d0.5_rel | -2762 | 5540.2 | 0 | -652 | 1319.63 | 0.96 |
| | | d0.6_rel | -2762 | 5540.2 | 0 | -652 | 1319.65 | 0.98 |
| | | d0.7_rel | -2762 | 5540.2 | 0 | -652 | 1319.78 | 1.11 |
| | | d0.8_rel | -2762 | 5540.2 | 0 | -652 | 1319.93 | 1.25 |
| | | d0.9_rel | -2762 | 5540.2 | 0 | -652 | 1320.06 | 1.38 |
| | | d1_rel | -2762 | 5540.2 | 0 | -652 | 1320.18 | 1.51 |
| Weight | Mom + Helper | d1_rel | -7621 | 15258.09 | 3.74 | -2394 | 4803.7 | 3.3 |
| | | d0.9_rel | -7621 | 15258.24 | 3.89 | -2394 | 4803.7 | 3.3 |
| | | d0.8_rel | -7621 | 15258.4 | 4.05 | -2394 | 4803.71 | 3.31 |
| | | d0.7_rel | -7621 | 15258.56 | 4.2 | -2394 | 4803.72 | 3.31 |
| | | d0.6_rel | -7621 | 15258.71 | 4.36 | -2394 | 4803.73 | 3.32 |
| | | d0.5_rel | -7621 | 15258.85 | 4.5 | -2394 | 4803.74 | 3.34 |
| | | d0.4_rel | -7621 | 15258.99 | 4.64 | -2394 | 4803.76 | 3.36 |
| | | d0.3_rel | -7622 | 15259.11 | 4.76 | -2394 | 4803.78 | 3.38 |
| | | d0.2_rel | -7622 | 15259.22 | 4.87 | -2394 | 4803.81 | 3.41 |
| | | d0.1_rel | -7622 | 15259.31 | 4.96 | -2394 | 4803.84 | 3.44 |
| | | d0_rel | -7622 | 15259.39 | 5.04 | -2394 | 4803.87 | 3.47 |
| Tarsus | | d0_rel | -2761 | 5538.81 | 0 | -652 | 1319.27 | 0.44 |
| | | d0.1_rel | -2761 | 5538.81 | 0 | -652 | 1319.34 | 0.52 |
| | | d0.2_rel | -2761 | 5538.81 | 0 | -652 | 1319.38 | 0.55 |
| | | d0.3_rel | -2761 | 5538.81 | 0 | -652 | 1319.42 | 0.6 |
| | | d0.4_rel | -2761 | 5538.81 | 0 | -652 | 1319.5 | 0.68 |
| | | d0.5_rel | -2761 | 5538.81 | 0 | -652 | 1319.52 | 0.69 |
| | | d0.6_rel | -2761 | 5538.81 | 0 | -652 | 1319.65 | 0.82 |
| | | d0.7_rel | -2761 | 5538.81 | 0 | -652 | 1319.78 | 0.95 |
| | | d0.8_rel | -2761 | 5538.81 | 0 | -652 | 1319.9 | 1.07 |
| | | d0.9_rel | -2761 | 5538.81 | 0 | -652 | 1320.01 | 1.19 |
| | | d1_rel | -2761 | 5538.81 | 0 | -652 | 1320.12 | 1.29 |
| Weight | Mom + Dad + Helper | d1_rel | -7617 | 15253.79 | 2.95 | -2392 | 4804.55 | 0.62 |
| | | d0.9_rel | -7617 | 15253.95 | 3.11 | -2394 | 4807.2 | 3.27 |
| | | d0.8_rel | -7617 | 15254.12 | 3.27 | -2394 | 4807.2 | 3.27 |
| | | d0.7_rel | -7617 | 15254.29 | 3.44 | -2394 | 4807.21 | 3.28 |
| | | d0.6_rel | -7617 | 15254.46 | 3.61 | -2394 | 4807.21 | 3.29 |
| | | d0.5_rel | -7617 | 15254.63 | 3.78 | -2394 | 4807.22 | 3.3 |
| | | d0.4_rel | -7617 | 15254.79 | 3.95 | -2394 | 4807.24 | 3.31 |
| | | d0.3_rel | -7617 | 15254.95 | 4.11 | -2394 | 4807.26 | 3.33 |
| | | d0.2_rel | -7618 | 15255.1 | 4.25 | -2394 | 4807.29 | 3.36 |
| | | d0.1_rel | -7618 | 15255.23 | 4.39 | -2394 | 4807.31 | 3.38 |
| | | d0_rel | -7618 | 15255.36 | 4.51 | -2394 | 4807.34 | 3.41 |
| Tarsus | | d0_rel | -2761 | 5542.81 | 0 | -651 | 1323 | 0.59 |
| | | d0.1_rel | -2761 | 5542.81 | 0 | -652 | 1323.03 | 0.62 |

| | | | | | | | | |
|---|---|---|---|---|---|---|---|---|
| | | d0.2_rel | -2761 | 5542.81 | 0 | -652 | 1323.08 | 0.67 |
| | | d0.3_rel | -2761 | 5542.81 | 0 | -652 | 1323.16 | 0.75 |
| | | d0.4_rel | -2761 | 5542.81 | 0 | -652 | 1323.18 | 0.77 |
| | | d0.5_rel | -2761 | 5542.81 | 0 | -652 | 1323.24 | 0.83 |
| | | d0.6_rel | -2761 | 5542.81 | 0 | -652 | 1323.33 | 0.92 |
| | | d0.7_rel | -2761 | 5542.81 | 0 | -652 | 1323.47 | 1.06 |
| | | d0.8_rel | -2761 | 5542.81 | 0 | -652 | 1323.6 | 1.19 |
| | | d0.9_rel | -2761 | 5542.81 | 0 | -652 | 1323.72 | 1.31 |
| | | d1_rel | -2761 | 5542.81 | 0 | -652 | 1323.83 | 1.42 |

**Supplementary Table 4. Fixed effect estimates for all models.** M = mom, D = dad, H = helper. Significant terms ($p$ < 0.05 from Wald tests) are bolded and standard errors are shown in parentheses.

| Trait | Model | Intercept | Sex (M) | Hatching Date | Brood Size | Number of Helpers | Inbreeding Coefficient | Age Measured | Territory Area |
|---|---|---|---|---|---|---|---|---|---|
| Nestling Weight | DGE Only | **43.3 (0.429)** | **0.756 (0.224)** | 0.158 (0.152) | **-1.49 (0.149)** | **0.752 (0.157)** | **-0.464 (0.159)** | – | 0.266 (0.165) |
| | M | **43.3 (0.428)** | **0.753 (0.224)** | 0.151 (0.152) | **-1.49 (0.149)** | **0.739 (0.158)** | **-0.46 (0.161)** | – | 0.268 (0.166) |
| | D | **43.2 (0.428)** | **0.744 (0.224)** | 0.155 (0.151) | **-1.5 (0.148)** | **0.697 (0.158)** | **-0.485 (0.16)** | – | 0.273 (0.168) |
| | H | **43.2 (0.431)** | **0.745 (0.224)** | 0.142 (0.151) | **-1.5 (0.148)** | **0.654 (0.189)** | **-0.486 (0.158)** | – | 0.278 (0.165) |
| | M+D | **43.2 (0.426)** | **0.743 (0.224)** | 0.151 (0.151) | **-1.51 (0.148)** | **0.69 (0.158)** | **-0.478 (0.161)** | – | 0.273 (0.169) |
| | M+H | **43.2 (0.43)** | **0.742 (0.224)** | 0.136 (0.151) | **-1.5 (0.148)** | **0.647 (0.189)** | **-0.482 (0.16)** | – | 0.279 (0.166) |
| | D+H | **43.2 (0.431)** | **0.733 (0.223)** | 0.139 (0.15) | **-1.51 (0.148)** | **0.613 (0.188)** | **-0.501 (0.159)** | – | 0.282 (0.168) |
| | M+D+H | **43.2 (0.429)** | **0.73 (0.223)** | 0.135 (0.15) | **-1.51 (0.148)** | **0.605 (0.189)** | **-0.495 (0.16)** | – | 0.285 (0.169) |
| Juvenile Weight | DGE Only | **67.6 (0.343)** | **4.12 (0.179)** | -0.0666 (0.123) | **-0.294 (0.116)** | -0.252 (0.123) | -0.112 (0.121) | 0.235 (0.142) | **0.403 (0.13)** |
| | M | **67.6 (0.343)** | **4.12 (0.179)** | -0.0582 (0.123) | **-0.291 (0.116)** | -0.248 (0.124) | -0.112 (0.122) | 0.242 (0.142) | **0.401 (0.13)** |
| | D | **67.6 (0.345)** | **4.11 (0.179)** | -0.05 (0.123) | **-0.292 (0.116)** | -0.248 (0.124) | -0.1 (0.122) | 0.236 (0.143) | **0.396 (0.13)** |
| | H | **67.6 (0.345)** | **4.12 (0.179)** | -0.0726 (0.123) | **-0.289 (0.116)** | -0.274 (0.144) | -0.105 (0.12) | 0.251 (0.142) | **0.395 (0.129)** |
| | M+D | **67.6 (0.345)** | **4.11 (0.179)** | -0.0484 (0.123) | **-0.292 (0.116)** | -0.246 (0.124) | -0.101 (0.122) | 0.238 (0.143) | **0.396 (0.131)** |
| | M+H | **67.6 (0.345)** | **4.12 (0.179)** | -0.0694 (0.123) | **-0.288 (0.116)** | -0.273 (0.144) | -0.105 (0.12) | 0.253 (0.142) | **0.394 (0.129)** |
| | D+H | **67.6 (0.346)** | **4.12 (0.179)** | -0.0594 (0.123) | **-0.288 (0.116)** | -0.268 (0.143) | -0.0949 (0.121) | 0.251 (0.142) | **0.39 (0.13)** |
| | M+D+H | **67.6 (0.346)** | **4.12 (0.179)** | -0.0594 (0.123) | **-0.288 (0.116)** | -0.268 (0.143) | -0.0949 (0.121) | 0.251 (0.142) | **0.39 (0.13)** |
| Nestling Tarsus | DGE Only | **29.2 (0.214)** | 0.13 (0.101) | **0.328 (0.0661)** | -0.0921 (0.065) | **0.173 (0.068)** | **-0.181 (0.0703)** | – | **0.156 (0.0704)** |
| | M | **29.2 (0.214)** | 0.126 (0.101) | **0.323 (0.066)** | -0.0941 (0.0651) | **0.169 (0.0681)** | **-0.178 (0.0711)** | – | **0.161 (0.0705)** |
| | D | **29.2 (0.214)** | 0.13 (0.101) | **0.328 (0.0661)** | -0.0925 (0.065) | **0.173 (0.0681)** | **-0.182 (0.0705)** | – | **0.156 (0.0705)** |
| | H | **29.2 (0.214)** | 0.13 (0.101) | **0.328 (0.0661)** | -0.0921 (0.065) | **0.173 (0.068)** | **-0.181 (0.0703)** | – | **0.156 (0.0704)** |
| | M+D | **29.2 (0.214)** | 0.126 (0.101) | **0.323 (0.066)** | -0.0941 (0.0651) | **0.169 (0.0681)** | **-0.178 (0.0711)** | – | **0.161 (0.0705)** |

| | | | | | | | | | |
|---|---|---|---|---|---|---|---|---|---|
| | M+H | **29.2 (0.214)** | 0.126 (0.101) | **0.323 (0.066)** | -0.0941 (0.0651) | **0.169 (0.0681)** | **-0.178 (0.0711)** | – | **0.161 (0.0705)** |
| | D+H | **29.2 (0.214)** | 0.13 (0.101) | **0.328 (0.0661)** | -0.0925 (0.065) | **0.173 (0.0681)** | **-0.182 (0.0705)** | – | **0.156 (0.0705)** |
| | M+D+H | **29.2 (0.214)** | 0.126 (0.101) | **0.323 (0.066)** | -0.0941 (0.0651) | **0.169 (0.0681)** | **-0.178 (0.0711)** | – | **0.161 (0.0705)** |
| Juvenile Tarsus | DGE Only | **35.8 (0.0867)** | **0.763 (0.0505)** | 0.0147 (0.0327) | **-0.118 (0.031)** | 0.0259 (0.0326) | -0.0185 (0.0324) | 0.0727 (0.0382) | 0.0365 (0.0343) |
| | M | **35.8 (0.0868)** | **0.762 (0.0504)** | 0.0174 (0.0327) | **-0.116 (0.031)** | 0.0251 (0.0326) | -0.0179 (0.0331) | 0.0732 (0.0383) | 0.0336 (0.0344) |
| | D | **35.8 (0.0873)** | **0.763 (0.0505)** | 0.0176 (0.0327) | **-0.117 (0.0311)** | 0.0279 (0.0326) | -0.0169 (0.0327) | 0.0741 (0.0383) | 0.0343 (0.0345) |
| | H | **35.8 (0.0872)** | **0.76 (0.0504)** | 0.0128 (0.0326) | **-0.119 (0.031)** | 0.0198 (0.0365) | -0.019 (0.0322) | 0.0722 (0.0382) | 0.0361 (0.0342) |
| | M+D | **35.8 (0.0872)** | **0.762 (0.0504)** | 0.0188 (0.0327) | **-0.116 (0.031)** | 0.0265 (0.0326) | -0.0173 (0.0332) | 0.0742 (0.0383) | 0.0329 (0.0346) |
| | M+H | **35.8 (0.0873)** | **0.759 (0.0504)** | 0.0153 (0.0326) | **-0.117 (0.031)** | 0.0192 (0.0365) | -0.0185 (0.0329) | 0.0723 (0.0382) | 0.033 (0.0344) |
| | D+H | **35.8 (0.0876)** | **0.759 (0.0504)** | 0.0164 (0.0326) | **-0.117 (0.0311)** | 0.0225 (0.037) | -0.0156 (0.0328) | 0.0724 (0.0382) | 0.0329 (0.0345) |
| | M+D+H | **35.8 (0.0876)** | **0.759 (0.0504)** | 0.017 (0.0326) | **-0.117 (0.031)** | 0.0214 (0.0369) | -0.0166 (0.0331) | 0.0725 (0.0382) | 0.0319 (0.0345) |
